# Recurrent deletions in the SARS-CoV-2 spike glycoprotein drive antibody escape

**DOI:** 10.1101/2020.11.19.389916

**Authors:** Kevin R. McCarthy, Linda J. Rennick, Sham Nambulli, Lindsey R. Robinson-McCarthy, William G. Bain, Ghady Haidar, W. Paul Duprex

## Abstract

Zoonotic pandemics, like that caused by SARS-CoV-2, can follow the spillover of animal viruses into highly susceptible human populations. Their descendants have adapted to the human host and evolved to evade immune pressure. Coronaviruses acquire substitutions more slowly than other RNA viruses, due to a proofreading polymerase. In the spike glycoprotein, we find recurrent deletions overcome this slow substitution rate. Deletion variants arise in diverse genetic and geographic backgrounds, transmit efficiently, and are present in novel lineages, including those of current global concern. They frequently occupy recurrent deletion regions (RDRs), which map to defined antibody epitopes. Deletions in RDRs confer resistance to neutralizing antibodies. By altering stretches of amino acids, deletions appear to accelerate SARS-CoV-2 antigenic evolution and may, more generally, drive adaptive evolution.

## Main text

SARS-CoV-2 emerged from a yet-to-be defined animal reservoir and initiated a pandemic in 2020 (*1–5*). It has acquired limited adaptions, most notably the D614G substitution in the spike (S) glycoprotein (*6–8*). Humoral immunity to S glycoprotein appears to be the strongest correlate of protection (*9*) and recently approved vaccines deliver this antigen by immunization. Coronaviruses like SARS-CoV-2 slowly acquire substitutions due to a proofreading RNA dependent RNA polymerase (RdRp) (*10, 11*). Other emerging respiratory viruses have produced pandemics followed by endemic human-to-human spread. The latter is often contingent upon the introduction of antigenic novelty that enables reinfection of previously immune individuals. Whether SARS-CoV-2 S glycoprotein will evolve altered antigenicity, or specifically how it may change in response to immune pressure, remains unknown. We and others have reported the acquisition of deletions in the amino (N)-terminal domain (NTD) of the S glycoprotein during long-term infections of often-immunocompromised patients (*12–15*). We have identified this as an evolutionary pattern defined by recurrent deletions that alter defined antibody epitopes. Unlike substitutions, deletions cannot be corrected by proofreading activity and this may accelerate adaptive evolution in SARS-CoV-2.

An immunocompromised cancer patient infected with SARS-CoV-2 was unable to clear the virus and succumbed to the infection 74 days after COVID-19 diagnosis (*15*). Treatment included Remdesivir, dexamethasone and two infusions of convalescent serum. We designate the individual as Pittsburgh long-term infection 1 (PLTI1). We consensus sequenced and cloned S genes directly from clinical material obtained 72 days following COVID-19 diagnosis and identified two variants with deletions in the NTD (Fig. 1A).

**Fig. 1.**
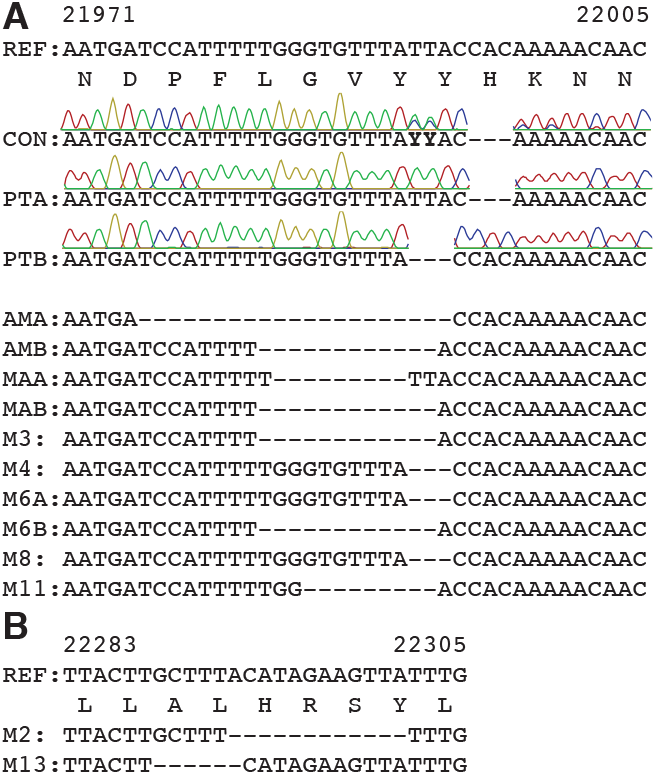
Deletions in SARS-CoV-2 spike arise during persistent infections of immunosuppressed patients. A. Top. Sequences of viruses isolated from PLTI1 (PT) and viruses from patients with deletions in the same NTD region. Chromatograms are shown for sequences from PLTI1, which include sequencing of bulk reverse transcription products (CON) and individual cDNA clones. Bottom. Sequences from other long term infections from individuals AM (18) MA-JL (MA) (19) and a MSK cohort (M) with individuals 2, 3, 4, 6, 8, 11, 13 (*13*) Letters (A&B) designate different variants from the same patient. (B) Sequences of viruses from two patients with deletions in a different region of the NTD. All sequences are aligned to reference sequence (REF) MN985325 (WA-1). Genetic analysis of patient isolates is in Fig. S1.

These data from PLTI1 and a similar report (*12*) prompted us to interrogate patient metadata sequences deposited in GISAID (*16*). In searching for similar viruses, we identified eight patients with deletions in the S glycoproteins of viruses sampled longitudinally over a period of weeks to months (Figs. 1A and S1A). For each, early time points had intact S sequences and later time points had deletions within the S gene. Six had deletions that were identical to, overlapping with, or adjacent to those in PLTI1. Deletions at a second site were present in viruses isolated from two other patients (Fig. 1B), reports on these patients have since been published (*13, 14*). Viruses from all but one patient could be distinguished from one another by nucleotide differences present at both early and late time points (Fig. S1B). On a tree of representative contemporaneously circulating isolates they form monophyletic clades making either a second community- or nosocomially-acquired infection unlikely (Fig. S1C). The most parsimonious explanation is these deletions arose independently due to a common selective pressure to produce strikingly convergent outcomes.

We searched the GISAID sequence database (*16*) for additional instances of deletions within S glycoproteins. From a dataset of 146,795 sequences (deposited from 12/01/2019 to 10/24/2020) we identified 1,108 viruses with deletions in the S gene. When mapped to the S gene, 90% occupied four discrete sites within the NTD (Fig. 2A). We term these important sites recurrent deletion regions (RDRs), numbering them 1-4 from the 5’ to 3’ end of the S gene. Deletions identified in patient samples correspond to RDR2 (Fig. 1A) and RDR4 (Fig. 1B). Most deletions appear to have arisen and been retained in replication competent viruses. Without selective pressure, in-frame deletions should occur one third of the time. However, we observed a preponderance of in-frame deletions with lengths of 3, 6, 9 and 12 (Fig. 2B). Among all deletions, 93% are in frame and do not produce a stop codon (Fig. 2C). In the NTD, >97% of deletions maintain the open reading frame. Other S glycoprotein domains do not follow this trend e.g. deletions in the receptor binding domain (RBD) and S2 preserve the reading frame 30% and 37% of the time, respectively.

**Fig. 2.**
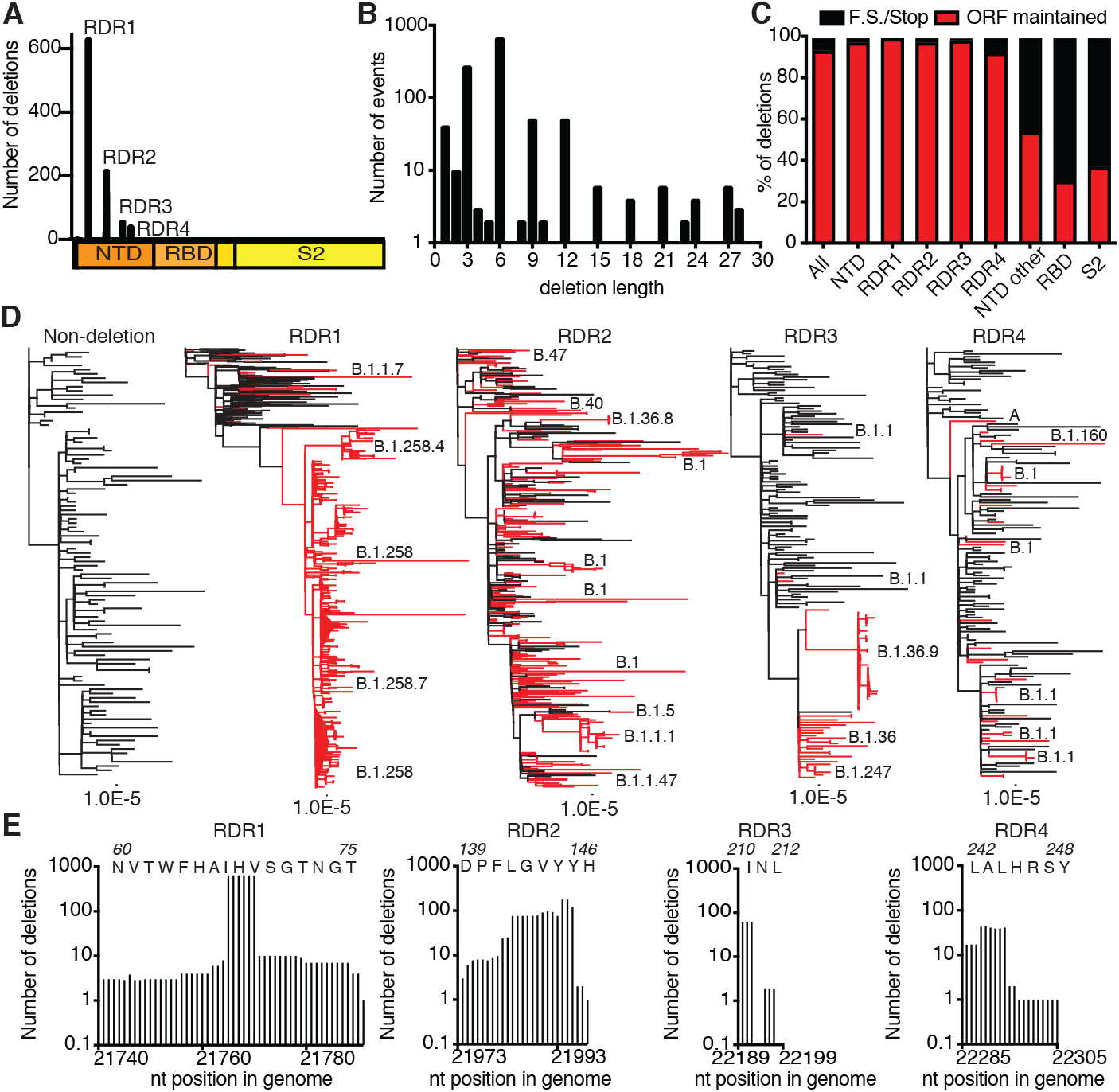
Identification and characterization of recurrent deletion regions in SARS-CoV-2 spike protein. A. Positional quantification of deleted nucleotides in S among GISAID sequences. We designate the four clusters recurrent deletion regions (RDRs)1-4.). B. Length distribution of deletions. C. The percentage of deletion events at the indicated site that either maintain the open reading frame or introduce a frameshift or premature stop codon (F.S./Stop). D. Phylogenetic analysis of deletion variants (red branches) and genetically diverse non-deletion variants (black branches). Specific deletion clades/lineages are identified. Maximum likelihood phylogenetic trees, rooted on NC_045512, were calculated with 1000 bootstrap replicates. Trees with branch labels are in Fig. S2. E. Abundance of nucleotide deletions in each RDR. Positions are defined by reference sequence MN985325, by codon (top) and nucleotide (below).

To trace the origins of RDR variants, we produced phylogenies for each with 101 additional genomes that sample much of the genetic diversity within the pandemic (Fig. 2D). The RDR variants interleave with non-deletion sequences and occupy distinct branches, indicating their recurrent generation. This is most pronounced for RDRs 1, 2 and 4 but also true of RDR 3, with conservatively four independent instances. RDR variants form distinct lineages/branches, most prominently in RDR1 (lineage B.1.258) and suggest human-to-human transmission events. We verified, using sequences with sufficient metadata that explicitly differentiate individuals, the transmission of a variant within each RDR between people (Fig. S2).

We defined the RDRs based upon peaks in the spectrum of S glycoprotein deletions. Deletion lengths and positions vary within RDRs 1, 2 and 4 (Fig. 2E). Variation is greatest in RDRs 2 and 4 with the loss of S glycoprotein residues 144/145 (adjacent tyrosine codons) in RDR2 and 243-244 in RDR4 appearing to be favored. In contrast, the loss of residues 69-70 accounts for the vast majority of RDR1 deletions. Based upon our phylogenetic analysis and supported by accompanying lineage classifications this two amino acid deletion has arisen independently at least thirteen times. RDR3 largely consists of three nucleotide (nt) deletions in codon 220.

We evaluated the genetic, geographic and temporal sampling of RDR variants (Fig. 3A-B). This analysis is limited to sequences deposited in GISAID (*16*) where sequences from specific nations and regions are overrepresented e.g. United Kingdom and other European countries. We show the distribution of all sequences within the database for reference. For RDRs 2 and 4 the genetic and geographic distributions largely mirror those of reported sequences. Variants of RDRs 1 and 3 are strongly polarized to specific clades and geographies. This is likely the result of successful lineages, circulating in regions with strong sequencing initiatives. Our temporal analysis indicates that RDR variants have been present throughout the pandemic (Fig. 3C). Specific variant lineages like B.1.258 (Fig. 2D) harboring Δ69-70 in RDR1 have rapidly risen to notable abundance (Fig. 3D). Circulation of B.1.36 with RDR3 Δ210 accounts for most of the RDR3 examples (Figs. 2D and 3 C&D). The abundance of RDR2 Δ144/145 is explained by independent deletion events followed by transmission (Figs. 2D and 3 C&D).

**Fig. 3.**
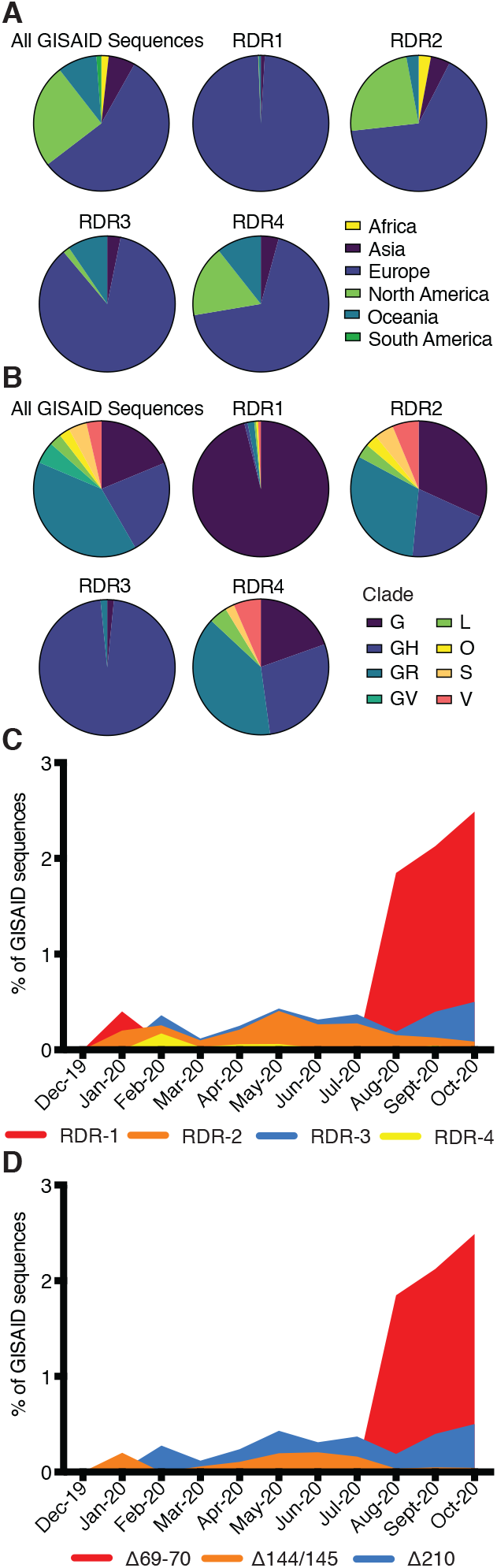
Geographic, genetic, and temporal abundance of RDR variants. Geographic (A) and genetic (B) distributions of RDR variants compared to the GISAID database (sequences from 12-1-2019 to 10-24-2020). GISAID clade classifications are used in B. C. Frequency of RDR variants among all complete genomes deposited in GISAID. D. Frequency of specific RDR deletion variants (numbered according to spike amino acids) among all GISAID variants. The plot of RDR3/Δ210 has been adjusted by 0.02 units on the Y-axis for visualization in panel C due to its overlap with RDR2 and this adjustment has been retained in panel D to make direct comparisons between panels.

The recurrence and convergence of RDR deletions, particularly during long-term infections, is indicative of adaptation in response to a common selective pressure. RDRs 2 and 4 and RDRs 1 and 3 occupy two distinct surfaces on the S glycoprotein NTD (Fig. 4A). Both sites contain antibody epitopes (*17–19*). The epitope for neutralizing antibody 4A8 is formed entirely by the beta sheets and extended connecting loops that harbor RDRs 2 and 4 (*17*). We generated a panel of S glycoprotein mutants representing the four RDRs to assess the impact deletions have on expression and antibody binding, we included an additional double mutant containing the deletions present in the B.1.1.7 variant of concern flagged initially in the United Kingdom. Cells were transfected with plasmids expressing these mutant glycoproteins and indirect immunofluorescence was used to determine if RDR deletions modulated 4A8 binding (Fig. 4B). Deletions at RDRs 1 and 3 had no impact on the binding of the monoclonal antibody, confirming that they alter independent sites. The three RDR2 deletions, the one RDR4 deletion and the double RDR1/2 deletions completely abolished binding of 4A8 whilst still allowing recognition by a monoclonal antibody targeting the RBD (Fig. 4B). Thus, convergent evolution operates in individual RDRs and between RDRs, exemplified by the same phenotype produced by deletions in RDR2 or RDR4.

**Fig. 4.**
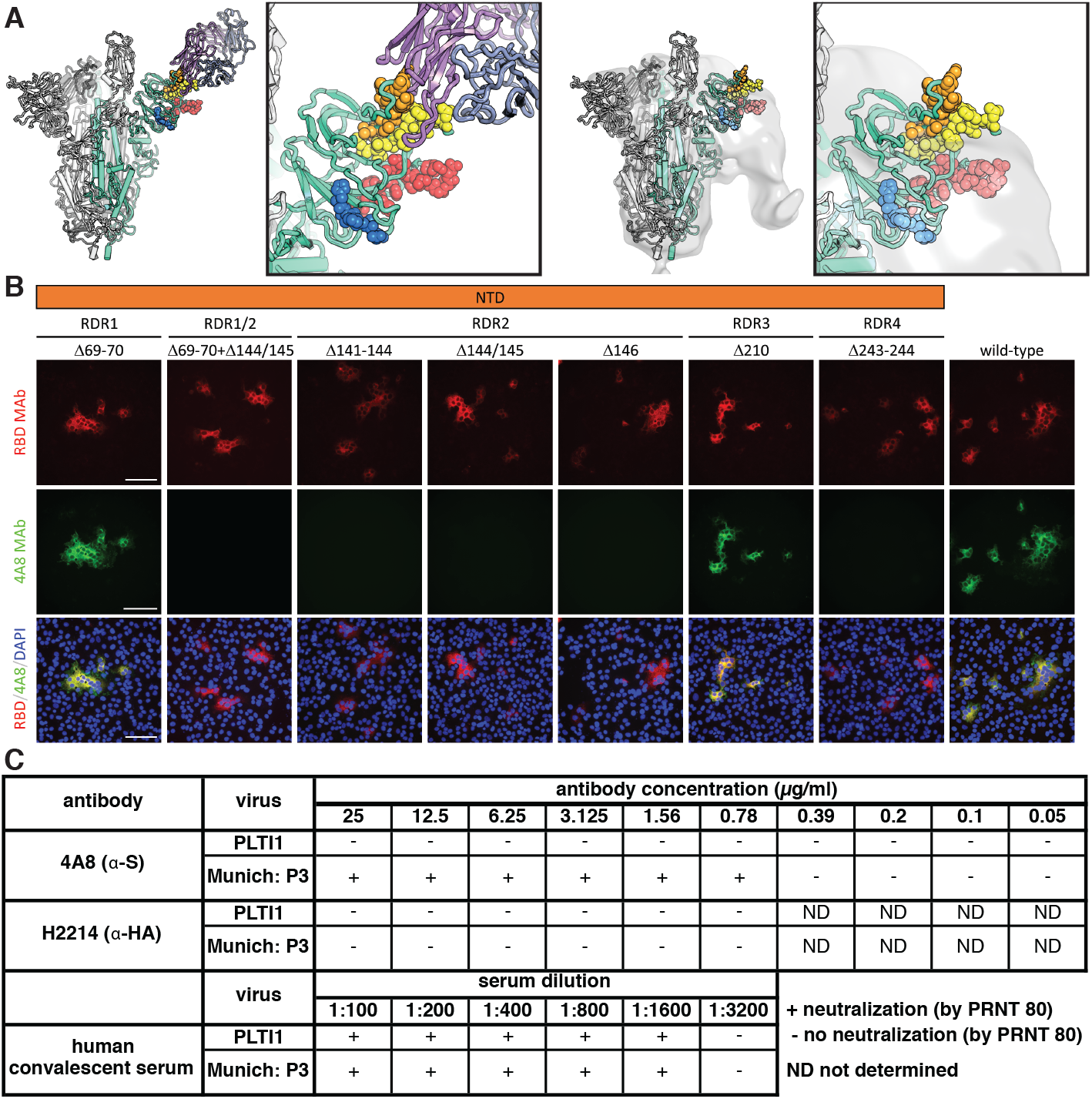
Deletions in the spike NTD alter its antigenicity. RDRs map to defined antigenic sites. (A) Top: A structure of antibody 4A8 (*17*) (PDB: 7C21) (purples) bound to one protomer (green) of a SARS-CoV-2 spike trimer (grays). RDRs 1-4 are colored red, orange, blue, and yellow, respectively, and shown in spheres. The interaction site is shown at right. Bottom: The electron microscopy density of COV57 serum Fabs (*18*) (EMDB emd_22125) fit to SARS-CoV-2 S glycoprotein trimer (PDB: 7C21). The same view of the interaction site is provided at right. (B) S glycoprotein distribution in Vero E6 cells at 24 h post-transfection with S protein deletion mutants, visualized by immunodetection in permeabilized cells. A monoclonal antibody against SARS-CoV-2 S protein receptor-binding domain (RBD MAb; red) detects all mutant forms of the protein (Δ69-70, Δ69-70+Δ141-144, Δ141-144, Δ144/145, Δ146, Δ210 and Δ243-244) and the unmodified protein (wild-type). 4A8 monoclonal antibody (4A8 MAb; green) does not detect mutants containing deletions in RDR2 or RDR4 (Δ69-70+Δ141-144, Δ141-144, Δ144/145, Δ146 and Δ243-244). Overlay images (RBD/4A8/DAPI) depict co-localization of the antibodies; nuclei were counterstained with DAPI (blue). The scale bars represent 100 μm. (C) Virus isolated from PLTI1 resists neutralization by 4A8. A non-deletion variant (Munich) is neutralized by 4A8, both are neutralized by convalescent serum and neither is neutralized by an influenza hemagglutinin binding antibody H2214 (*29*).

We assayed whether RDR variants escape the activity of a neutralizing antibody using the non-plaque purified viral population from PLTI1. This viral stock was completely resistant to neutralization by 4A8, while an isolate with authentic RDRs (*20*) was neutralized (Fig. 4C). We used a high titer neutralizing human convalescent polyclonal antiserum to demonstrate that both viral stocks could be neutralized efficiently. These data demonstrate that naturally arising and circulating variants of SARS-CoV-2 have altered antigenicity. We used a range of high, medium and low titer neutralizing human convalescent polyclonal antisera to assess if there was an appreciable difference in neutralization between the S glycoprotein-deleted and undeleted viruses. No major difference was observed suggesting that many more changes would be required to generate serologically distinct SARS-CoV-2 variants (Table S1).

Coronaviruses, including SARS-CoV-2, have lower substitution rates than other RNA viruses due to an RdRp with proofreading activity (*10, 11*). However, proofreading cannot correct deletions. We find that adaptive evolution of S glycoprotein is augmented by a tolerance for deletions, particularly within RDRs. The RDRs occupy defined antibody epitopes within the NTD (*17–19*) and deletions at multiple sites confer resistance to a neutralizing antibody. Deletions represent a generalizable mechanism through which S glycoprotein rapidly acquires genetic and antigenic novelty of SARS-CoV-2.

Fitness of RDR variants is evident by their representation in the consensus genomes from patients, transmission between individuals and presence in emergent lineages. Initially documented in the context of long-term infections of immunosuppressed patients, specific variants transmit efficiently between immunocompetent individuals. Characterization of unique cases led to the very early identification of RDR variants that are escape mutants. Since deletions are a product of replication, they will occur at a certain rate and variants are likely to emerge in otherwise healthy populations. Indeed, influenza explores variation that approximates future antigenic drift in immunosuppressed patients (*21*).

The RDRs occupy defined antibody epitopes within the S glycoprotein NTD. Selected *in vivo*, these deletion variants resist neutralization by monoclonal antibodies. Viruses cultured *in vitro* in the presence of immune serum have also acquired substitutions in RDR2 that confer neutralization resistance (*22*). Potent neutralizing responses and an array of monoclonal antibodies are directed to the RBD (*18, 19, 23*). A growing number of NTD directed antibodies have been identified (*24, 25*). Why antibody escape in nature is most evident in the NTD highlights a discrepancy and this requires further study.

During evaluation of this manuscript, RDR variants have been associated with numerous lineages of global concern. RDR variants independently emerged in farmed mink (Cluster 5) initiating culls and regional lockdowns (*26*). The recently identified B.1.1.7 (*27*) and B.1.351 (*28*), first reported in the United Kingdom and South Africa, have deletions in RDRs 1/2 and 4, respectively. Notably the RDR 2 and 4 deletions are functionally convergent, modifying the same antibody epitope conferring neutralization resistance. Additional circulating RDR variants have gone virtually unnoticed, while RBD substitutions receive considerable attention. Given the rate of substitution and the scale of the pandemic these mutations are repeatedly sampled in SARS-CoV-2 infected individuals daily. Success of SARS-CoV-2 lineages is likely dependent upon their genetic context (including deletions) and circumstance in which they emerge. Efforts to track and monitor RDR variants are vital.

## Materials and Methods

### Determination of PLTI1 patient spike gene sequences

To determine the consensus sequence of SARS-CoV-2 S in the patient endotracheal aspirate sample collected at day 72 (*15*), RNA was isolated from the sample using TRIzol LS (Thermo Fisher Scientific), cDNA was generated using the Superscript III first strand synthesis system (Thermo Fisher Scientific) and random hexamers, DNA was amplified using Phusion DNA polymerase (New England BioLabs) and SARS-CoV-2 specific primers surrounding the open reading frame for the spike protein, and the consensus sequence was determined by Sanger sequencing (Genewiz) using SARS-CoV-2 specific primers. The amplified DNA product was also cloned into pCR Blunt II TOPO vector using a Zero Blunt TOPO PCR Cloning Kit (Thermo Fisher Scientific) and the spike NTD sequence of individual clones was determined by Sanger sequencing (Genewiz) using M13F and M13R primers. Individual clone sequences are available with accession numbers MW269404 and MW269555.

### Sequence analysis

Sequences were obtained from the publically available GISAID database (*16*) and acknowledged in supporting Table 1. Our dataset was composed of SARS-CoV-2 sequences collected and deposited between 12-1-19 and 10-24-20. Sequence analysis was performed in Geneious (Biomatters, New Zealand). To identify deletion variants in S gene, sequences were mapped to NCBI reference sequence MN985325 (SARS-CoV-2/human/USA/WA-CDC-WA1/2020), the S gene open reading frame was extracted, remapped to reference and parsed for deletions using a search for gaps function. Sequences with deletions were manually extracted for subsequent analysis.

All identified deletion and non-deletion variants were aligned in MAFFT (*30, 31*) and adjusted manually in recurrent deletion regions for consistency. To evaluate the phylogenetic relationships of the long-term infections with reference to a sample of genetic diversity of contemporaneously circulating isolates, we used sequences from New York (where most patients were treated) from a time that most patients had their earliest reported samples, mid March through April. Reference sequences NC_045512 and MN985325 were included. These and long-term patient sequences were aligned using MAFFT (*30, 31*). FastTree (*32*) was used to generate a preliminary phylogeny, which we used to identify representatives of most clades and those sequences that interleaved between patients. The final tree, using this subset of sequences was produced using RAxML (*33*). To place RDR variants within a representative sample of genetic diversity we identified two high quality representatives without deletions from each lineage from which we identified a deletion variant. We attempted to find one temporarily early and late sequence when able to do so. For RDR transmission in individual nations, phylogenetic analyses utilized all sequences in our dataset from a country at a specific time, or in the case of Senegalese sequences the entirety of the pandemic. For non-Senegalese samples, sequences obtained within 1-2 months of the variants of interest were aligned to MN985325 using MAFFT (*30, 31*). FastTree (*32*) was used to generate a preliminary phylogeny from which we extracted the sequences corresponding to the lineage of interest and adjacent outgroups. These sequences were realigned using MAFFT. Maximum- Likelihood phylogenetic trees were calculated using RAxML (*33*) using a general time reversible model with optimization of substitution rates (GTR GAMMA setting), starting with a completely random tree, using rapid Bootstrapping and search for best-scoring ML tree. Between 1,000 and 10,000 bootstraps of support were performed.

### Cell lines

Human 293F cells were maintained at 37° Celsius with 5% CO_2_ in FreeStyle 293 Expression Medium (ThermoFisher) supplemented with penicillin and streptomycin. Vero E6 cells were maintained at 37° Celsius with 5% CO_2_ in high glucose DMEM (Invitrogen) supplemented with 1% (v/v) Glutamax (Invitrogen) and 10% (v/v) fetal bovine serum (Invitrogen).

### Recombinant IgG expression and purification

The heavy and light chain variable domains of 4A8 (*17*) was synthesized by Integrated DNA Technologies (Coralville, Iowa) and cloned into a modified human pVRC8400 expression vector encoding for full length human IgG1 heavy chains and human kappa light chains. Plasmids encoding influenza hemagglutinin-specific antibody H2214 have been described previously (*29*). IgGs were produced by polyethylenimine (PEI) facilitated, transient transfection of 293F cells that were maintained in FreeStyle 293 Expression Medium. Transfection complexes were prepared in Opti-MEM and added to cells. Five days post-transfection (d.p.t.) supernatants were harvested, clarified by low-speed centrifugation, adjusted to pH 5 by addition of 1 M 2-(N-morpholino)ethanesulfonic acid (MES) (pH 5.0), and incubated overnight with Pierce Protein G Agarose resin (Pierce, ThermoFisher). The resin was collected in a chromatography column, washed with a column volume of 100 mM sodium chloride 20 mM (MES) (pH 5.0) and eluted in 0.1 M glycine (pH 2.5) which was immediately neutralized by 1 M TRIS(hydroxymethyl)aminomethane (pH 8). IgGs were then dialyzed against phosphate buffered saline (PBS) pH 7.4.

### Cloning and transfection of SARS-CoV-2 spike protein deletion mutants

A series of deletion mutants were generated in HDM_SARS2_Spike_del21_D614G (*34*) a plasmid containing SARS-CoV-2 S protein lacking the 21 C-terminal amino acids. HDM_SARS2_Spike_del21_D614G was a gift from Jesse Bloom (Addgene plasmid # 158762; n2t.net/addgene:158762; RRID:Addgene_158762). Cloning strategies were designed to delete S protein amino acids 69-70 (Δ69-70), 141-144 (Δ141-144), 144/145 (Δ144/145), 146 (Δ146), 210 (Δ210), 243-244 (Δ243-244) or 69-70 and 144/145 (Δ69-70+Δ144/145). Appropriate gBlocks were generated synthetically (Integrated DNA Technologies) and cloned into HDM_SARS2_Spike_del21_D614G by Gibson Assembly using NEBuilder HiFi DNA Assembly Master Mix (New England Biolabs). Assemblies were transformed into DH5-alpha chemically competent cells (New England Biolabs) and correct clones were identified by restriction profile and Sanger sequencing (Genewiz) of small scale plasmid preparations from individual bacterial clones. Plasmid DNA for transfections was prepared using a HiSpeed Plasmid Midi Kit (Qiagen). Vero E6 cells were seeded into 24 well trays at 10^5^ cells per well. After overnight incubation at 37° Celsius, 5% (v/v) CO_2_, the cells were rinsed with Opti-MEM (Invitrogen), 1ml/well Opti-MEM was added and cells were incubated at 37° Celsius, 5% (v/v) CO_2_ for 30 minutes. Transfection mixes were prepared, according to manufacturer’s instructions, containing 200 ng/well of plasmid DNA with 3 μl per μg DNA of Lipofectamine 2000 (Invitrogen). After the 30 minute incubation Opti-MEM in the wells was replaced with 500 μl per well Opti-MEM and 100 μl per well of transfection mixes were added. Transfected cells were incubated at 37° Celsius, 5% (v/v) CO_2_ for 24 hours.

### Indirect immunofluorescence assay

Indirect immunofluorescence was performed as previously reported (*20*). Briefly, cells transfected with the SARS-CoV-2 S protein deletion mutants and controls were washed once with DPBS (Fisher Scientific), fixed with 4% (w/v) paraformaldehyde in PBS (Boston Bioproducts) for 20 minutes at room temperature, rinsed twice with DPBS and permeabilized with 0.1% (v/v) Triton-X100 (Sigma) in DPBS for 30 minutes at 37° Celsius. Primary antibodies [rabbit anti-SARS-CoV-2 S monoclonal antibody, 40150-R007, Sino Biological, 1/700 dilution and human 4A8 monoclonal antibody, 1 μg/ml, in PBS containing 0.1% (v/v) Triton X-100] were added and incubated at 37° Celsius for 1 hour. Cells were washed three times with DPBS and secondary antibodies [goat anti-rabbit Alexa Fluor-568, Invitrogen, and goat anti-human Alexa Fluor-488, Invitrogen, diluted 1:400 in DPBS containing 0.1% (v/v) Triton X-100 were added and incubated at 37° Celsius for 1 hour. Cells were washed three times with DPBS and nuclei were counterstained with 4’,6-diamidino-2-phenylindole (DAPI) nuclear stain (300 nM DAPI stain solution in PBS; Invitrogen) for 10 minutes at room temperature. Fluorescence was observed with a DMi 8 UV microscope (Leica) and photomicrographs were acquired using a camera (Leica) and LAS X software (Leica). Appropriate controls were included to determine antibody specificity.

### Virus neutralization assays

4A8 monoclonal antibody was diluted to 50 μg/ml in Opti-MEM which was used to prepare 2-fold serial dilutions to 0.1 μg/ml in Opti-MEM. An identical dilution series was prepared using H2214 monoclonal antibody as a negative control. Human convalescent sera samples were diluted in appropriate 2-fold series depending on their neutralization titers. Each antibody concentration or serum dilution (100 μl) was mixed with 100 μl of PLTI1 or Munich: P3 (*20*) viruses containing 50 plaque forming units (P.F.U.) of virus in Opti-MEM. These mixes were used in neutralization assays as previously described (*20*).

### Structure visualization

Structural figures were rendered in Pymol (The PyMOL Molecular Graphics System, Version 2.0 Schrödinger, LLC).

## Acknowledgments

We gratefully acknowledge the authors from the originating laboratories and the submitting laboratories, who generated and shared via GISAID genetic sequence data on which this research is based (Table S2). We thank Stephen C. Harrison for his support. We thank Dr. Alison Morris, Dr. Bryan McVerry, Dr. Georgios Kitsios, Dr. Barbara Methe, Heather Michael, Michelle Busch, John Ries, and Caitlin Schaefer at the University of Pittsburgh, as well as the physicians, nurses, and respiratory therapists at the University of Pittsburgh Medical Center Shadyside-Presbyterian Hospital intensive care units for assistance with collection and processing of the endotracheal aspirate sample.

## Author contribution

K.R.M., L.J.R., S.N., L.R.R.M. and W.P.D. designed the experiments. K.R.M., L.J.R., S.N. and L.R.R.M. performed the experiments. K.R.M., L.J.R., S.N., L.R.R.M. and W.P.D. analyzed data. W.G.B. and G.H. provided reagents and samples K.R.M., L.J.R., S.N., L.R.R.M. and W.P.D. wrote the manuscript.

## Data availability

Sequences from PLTI1 were deposited in NCBI GenBank under accession numbers MW269404 and MW269555. All other sequences are available via the GISAID SARS-CoV-2 sequence database (www.gsaid.org).

This work is licensed under a Creative Commons Attribution 4.0 International (CC BY 4.0) license, which permits unrestricted use, distribution, and reproduction in any medium, provided the original work is properly cited. To view a copy of this license, visit https://creativecommons.org/licenses/by/4.0/. This license does not apply to figures/photos/artwork or other content included in the article that is credited to a third party; obtain authorization from the rights holder before using such material.

