## Supporting Data for "Recurrent deletions in the SARS-CoV-2 spike glycoprotein drive antibody escape"

**A**

| MA-JL-D | | NY-MSK-2 | | NY-MSK-3 | | NY-MSK-4 | |
|---|---|---|---|---|---|---|---|
| Date | Accession | Date | Accession | Date | Accession | Date | Accession |
| 4/28/20 | EPI_ISL_593478 | 3/30/20 | EPI_ISL_583428 | 3/19/20 | EPI_ISL_583431 | 4/13/20 | EPI_ISL_583439 |
| 5/5/20 | EPI_ISL_593479 | 3/30/20 | EPI_ISL_583429 | 3/19/20 | EPI_ISL_583434 | 4/13/20 | EPI_ISL_583441 |
| 6/24/20 | EPI_ISL_593480 | 4/23/20 | EPI_ISL_583430 | 4/2/20 | EPI_ISL_583432 | 5/6/20 | EPI_ISL_583440 |
| 6/30/20 | EPI_ISL_593553 | | | 4/2/20 | EPI_ISL_583435 | 5/6/20 | EPI_ISL_583442 |
| 8/16/20 | EPI_ISL_593554 | | | 4/13/20 | EPI_ISL_583433 | 5/28/20 | EPI_ISL_583443 |
| 8/18/20 | EPI_ISL_593555 | | | 4/13/20 | EPI_ISL_583436 | | |
| 8/31/20 | EPI_ISL_593556 | | | 4/21/20 | EPI_ISL_583437 | | |
| 9/3/20 | EPI_ISL_593557 | | | 5/11/20 | EPI_ISL_583438 | | |
| 9/9/20 | EPI_ISL_593558 | | | | | | |

| NY-MSK-6 | | NY-MSK-8 | | NY-MSK-11 | | NY-MSK-13 | |
|---|---|---|---|---|---|---|---|
| Date | Accession | Date | Accession | Date | Accession | Date | Accession |
| 4/30/20 | EPI_ISL_583446 | 4/20/20 | EPI_ISL_583456 | 3/24/20 | EPI_ISL_583466 | 3/17/20 | EPI_ISL_583469 |
| 5/6/20 | EPI_ISL_583447 | 4/20/20 | EPI_ISL_583458 | 3/24/20 | EPI_ISL_583467 | 5/4/20 | EPI_ISL_583470 |
| 5/12/20 | EPI_ISL_583448 | 5/7/20 | EPI_ISL_583459 | 4/16/20 | EPI_ISL_583468 | 7/18/20 | EPI_ISL_583471 |
| 5/19/20 | EPI_ISL_583449 | 5/7/20 | EPI_ISL_583457 | | | | |
| 5/25/20 | EPI_ISL_583450 | 5/12/20 | EPI_ISL_583460 | | | | |
| 6/4/20 | EPI_ISL_583451 | | | | | | |
| 7/22/20 | EPI_ISL_583452 | | | | | | |
| 7/22/20 | EPI_ISL_583453 | | | | | | |
| 7/22/20 | EPI_ISL_583454 | | | | | | |

**B**

| Patient identifier | Substitution 1 | Substitution 2 | Substitution 3 | Substitution 4 | Substitution 5 | Substitution 6 | Substitution 7 |
|---|---|---|---|---|---|---|---|
| MA-JL-D | U1336C | G7936C | C16580T | C27881T | - | - | - |
| NY-MSK-2 | C5392T | C27236T | - | - | - | - | - |
| NY-MSK-3 | A2018G | G4790A | C24518T | - | - | - | - |
| NY-MSK-4 | A17376G | G28881A | G28882A | G28883C | - | - | - |
| NY-MSK-6 | G6358A | G11083T | C11916U | C18998U | U25233C | C25603T | G29540A |
| NY-MSK-8 | - | - | - | - | - | - | - |
| NY-MSK-11 | C19593U | - | - | - | - | - | - |
| NY-MSK-13 | C920U | - | - | - | - | - | - |

**C**

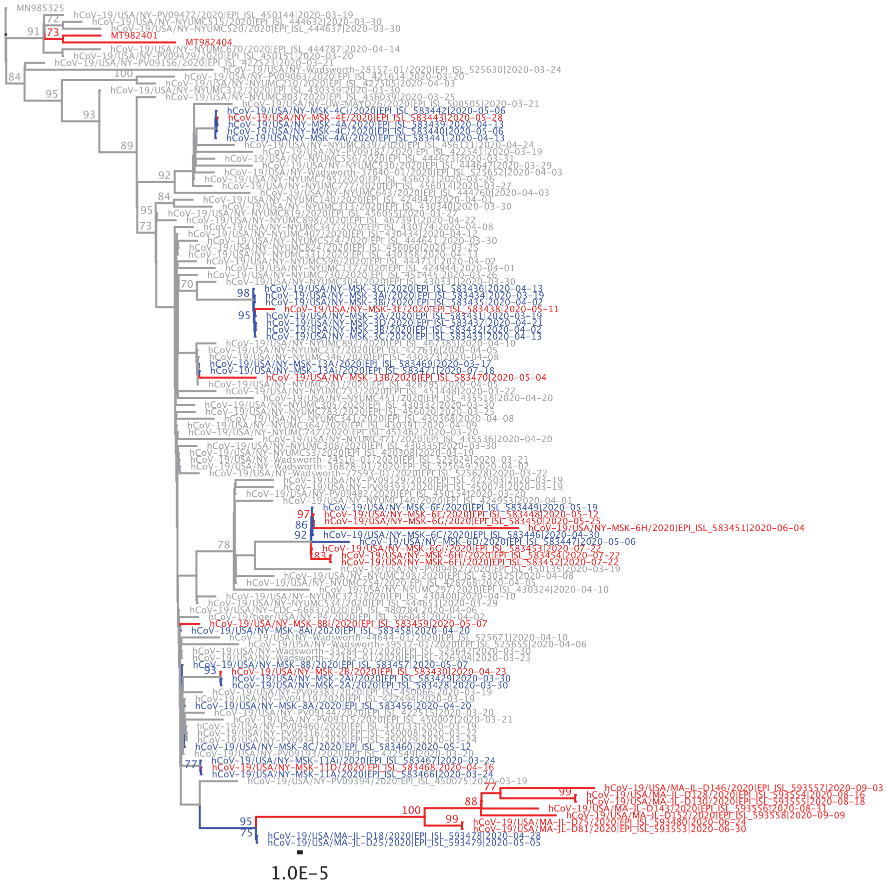

**Fig. S1.**

Information for the longitudinally sampled patients that were identified in the GISAID database and detailed in Figure 1. (A) Date of collection and GISAID accession number for each sequence. (B) Identifying substitutions unique to each patient among the longitudinally sampled patients reported here. (C) Phylogeny of RDR variants against a backdrop of contemporaneously circulating viruses. Maximum likelihood phylogenetic trees, rooted on NC_045512, were calculated with 1000 bootstrap replicates.

**A**

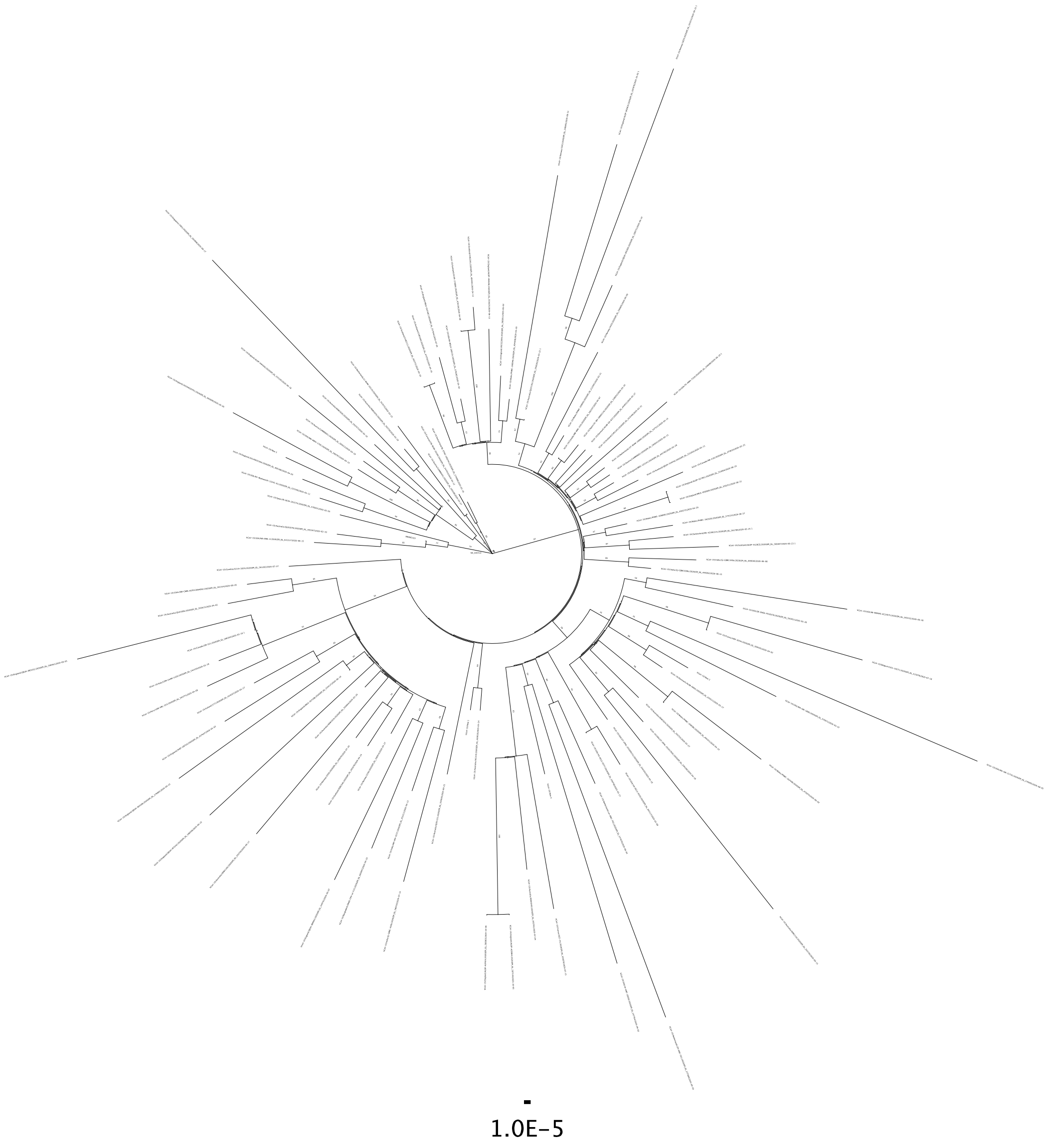

**Fig. S2.**

Phylogenetic analysis of RDR variants. Phylogenetic trees from Figure 2 with branch labels. (A) Non-deletion variants. (B). RDR1. (C) RDR2. (D) RDR3. (E) RDR4. Non-deletion variants are colored black and RDR variants are colored red.

**B**

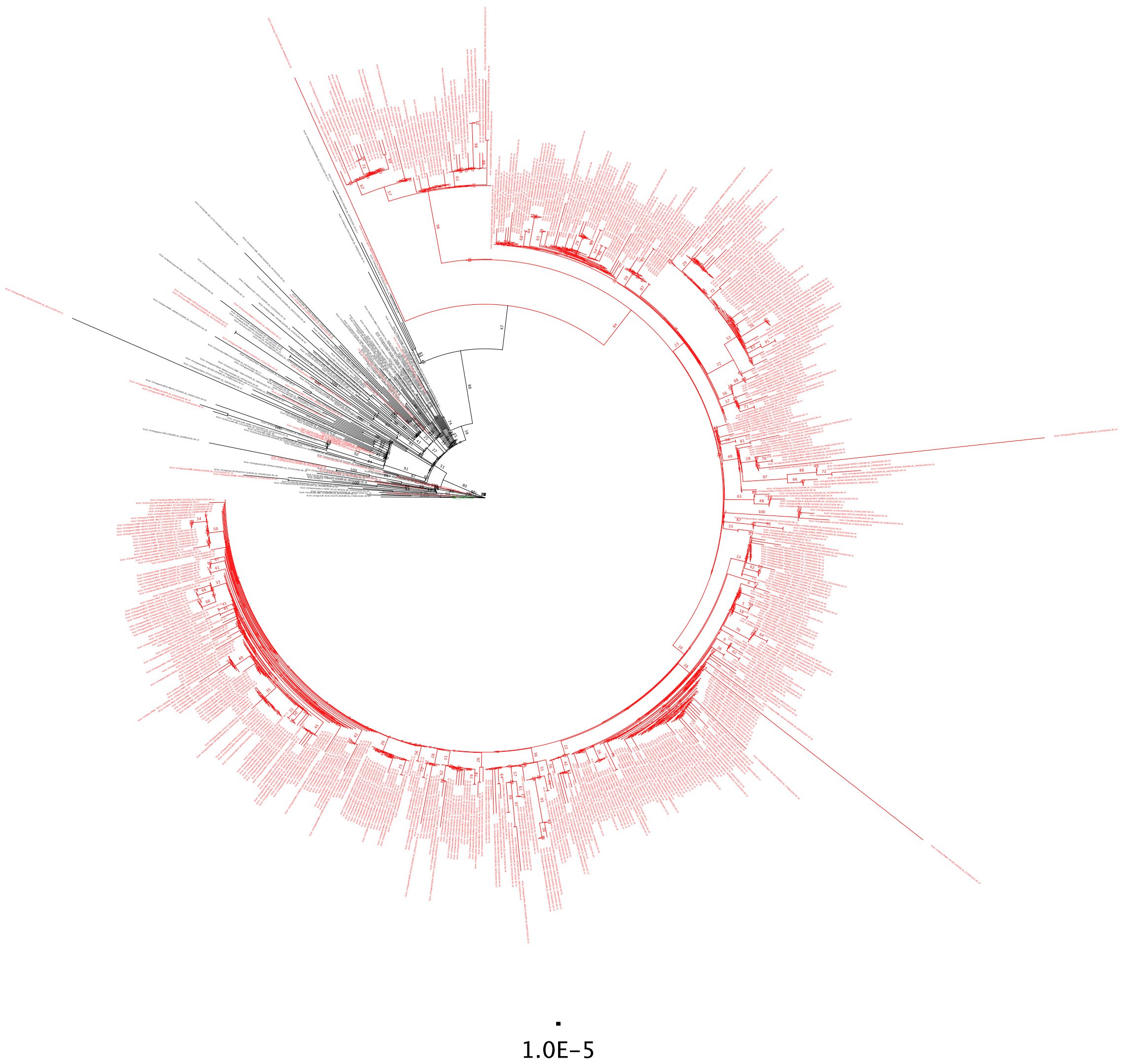


**Fig. S2.**
Continued.

**C**

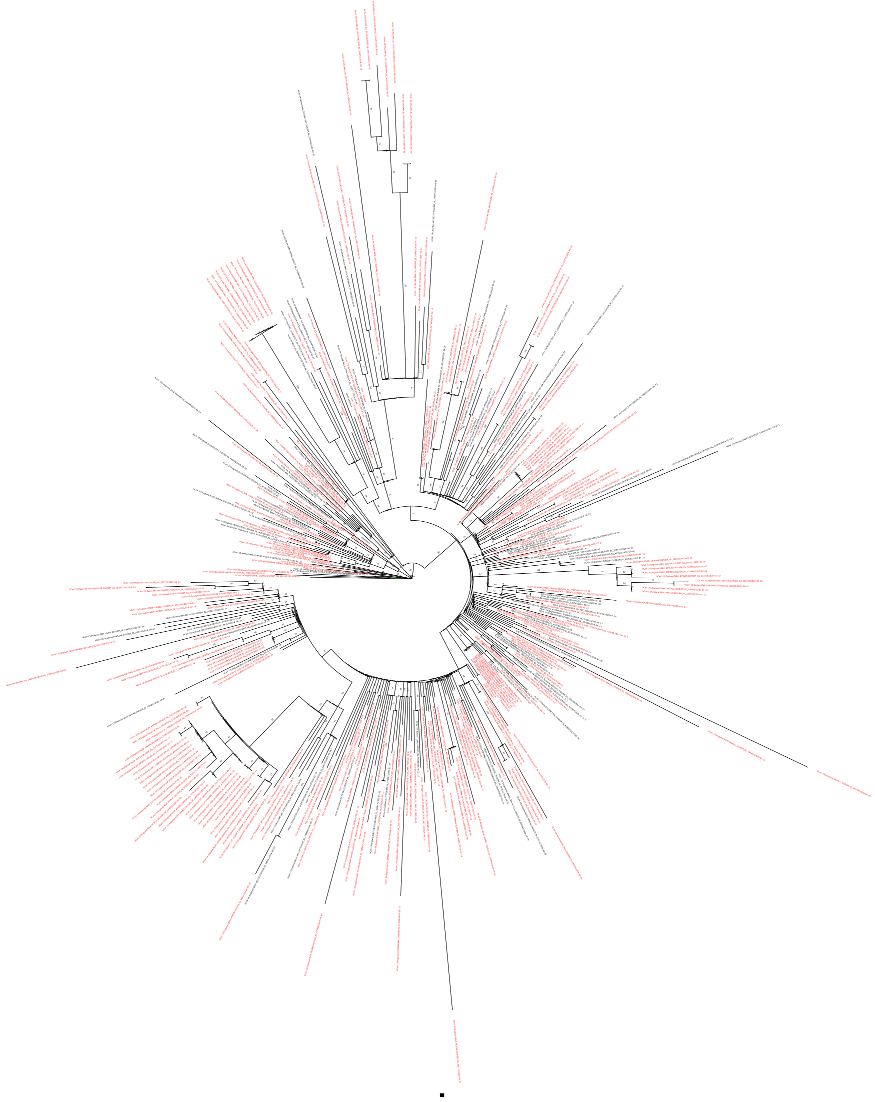


**Fig. S2.**
Continued

**D**

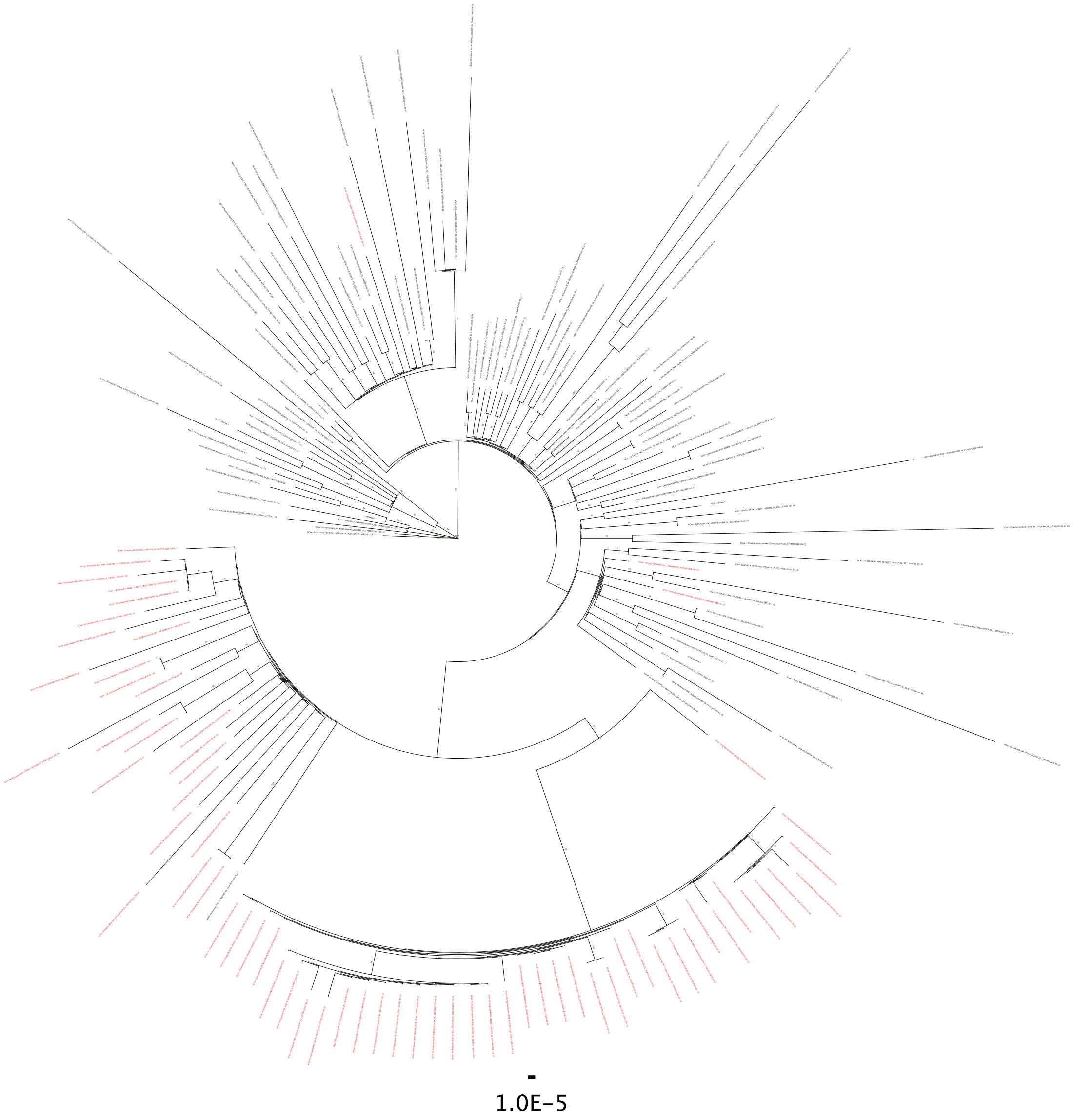


**Fig. S2.**
Continued.

E

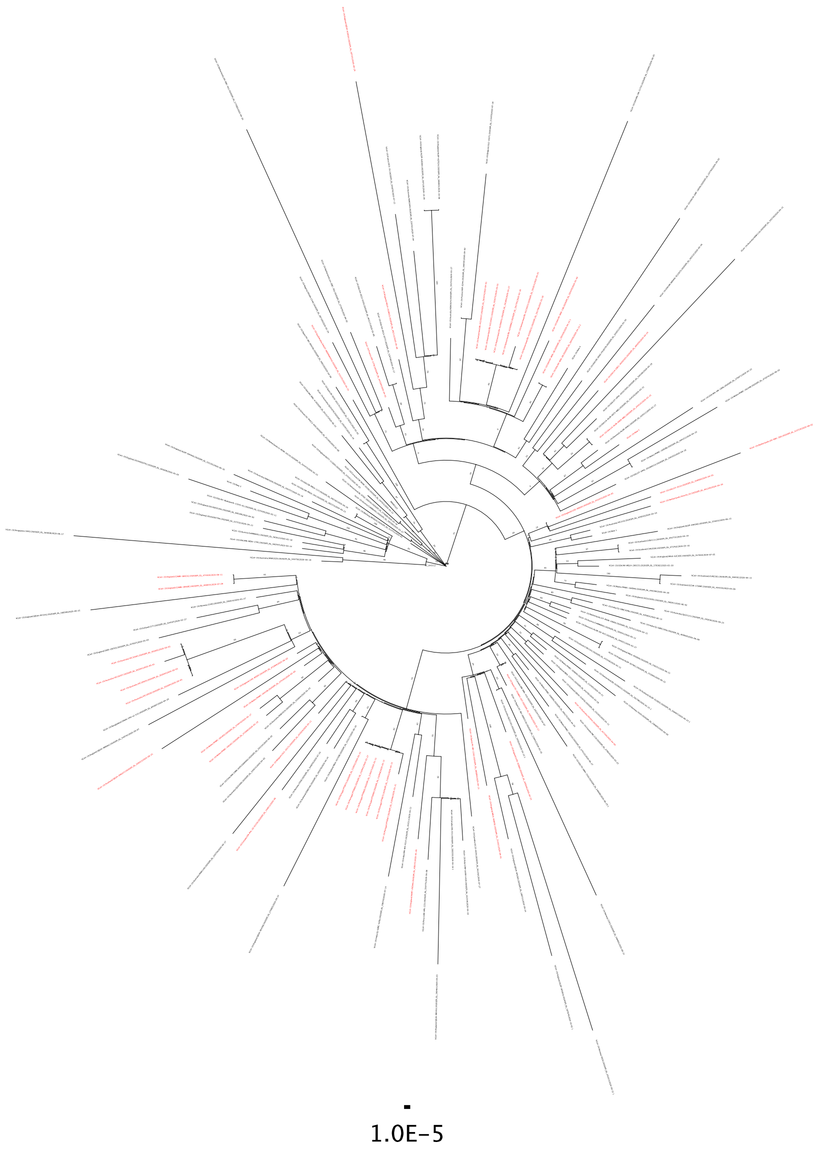


**Fig. S2.**
Continued.

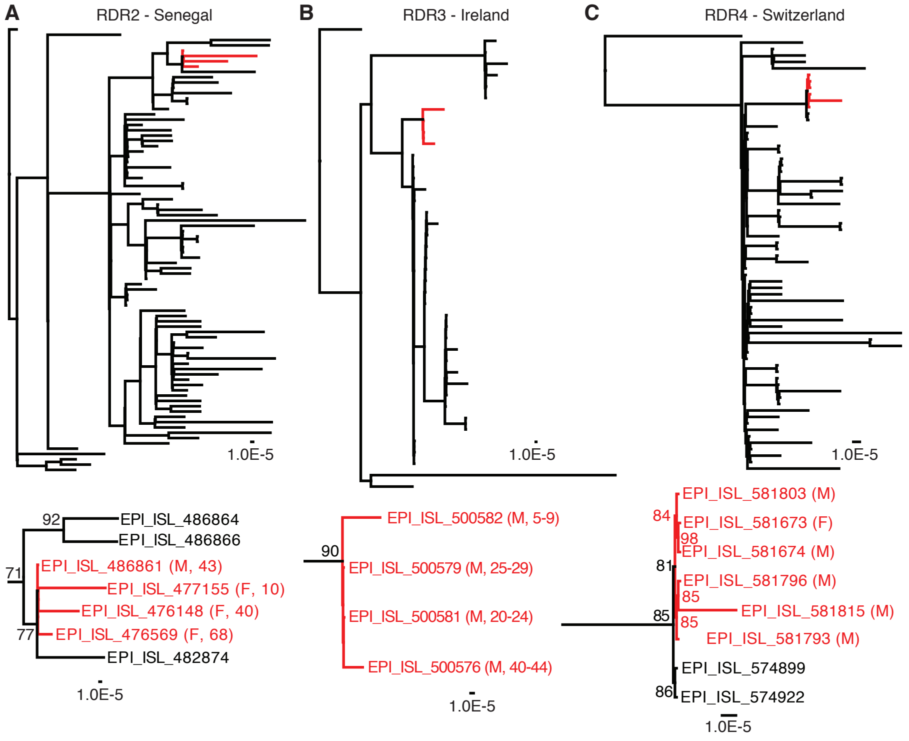


**Fig. S3**

Variants of all four RDRs transmit between humans. A-C Phylogenetic trees and patient data showing likely transmission of RDR variants between humans. We note two patients in France (male, age 58, sample collection 09-01-2020, EPI_ISL_582112 and female, age 59, sample collection 09-07-2020, EPI_ISL_582120) were found to have viruses that where 100% identical, including a six-nucleotide deletion in RDR1. A. Transmission of an RDR2 variant among 4 individuals in Senegal (deletion positions 21,991-21,994). B. Transmission cluster of an RDR3 variant (deletion positions 22,189-22,192) among four individuals in Ireland. C. Transmission of an RDR4 variant (deletion positions 22,281-22,290) among at least one male and female in Switzerland. Maximum likelihood phylogenetic trees are rooted on MN985325 and were calculated with 10,000 (A and B) or 1000 (C) bootstrap replicates. Branches with transmitted RDR variants are colored red and detailed below. Patient data differentiating individual patients is provided. (D-F). Full Phylogenetic analysis of transmitted RDR 2, 3, and 4 variants. For clarity all nodes with bootstrap values above 70 are labeled

**D**

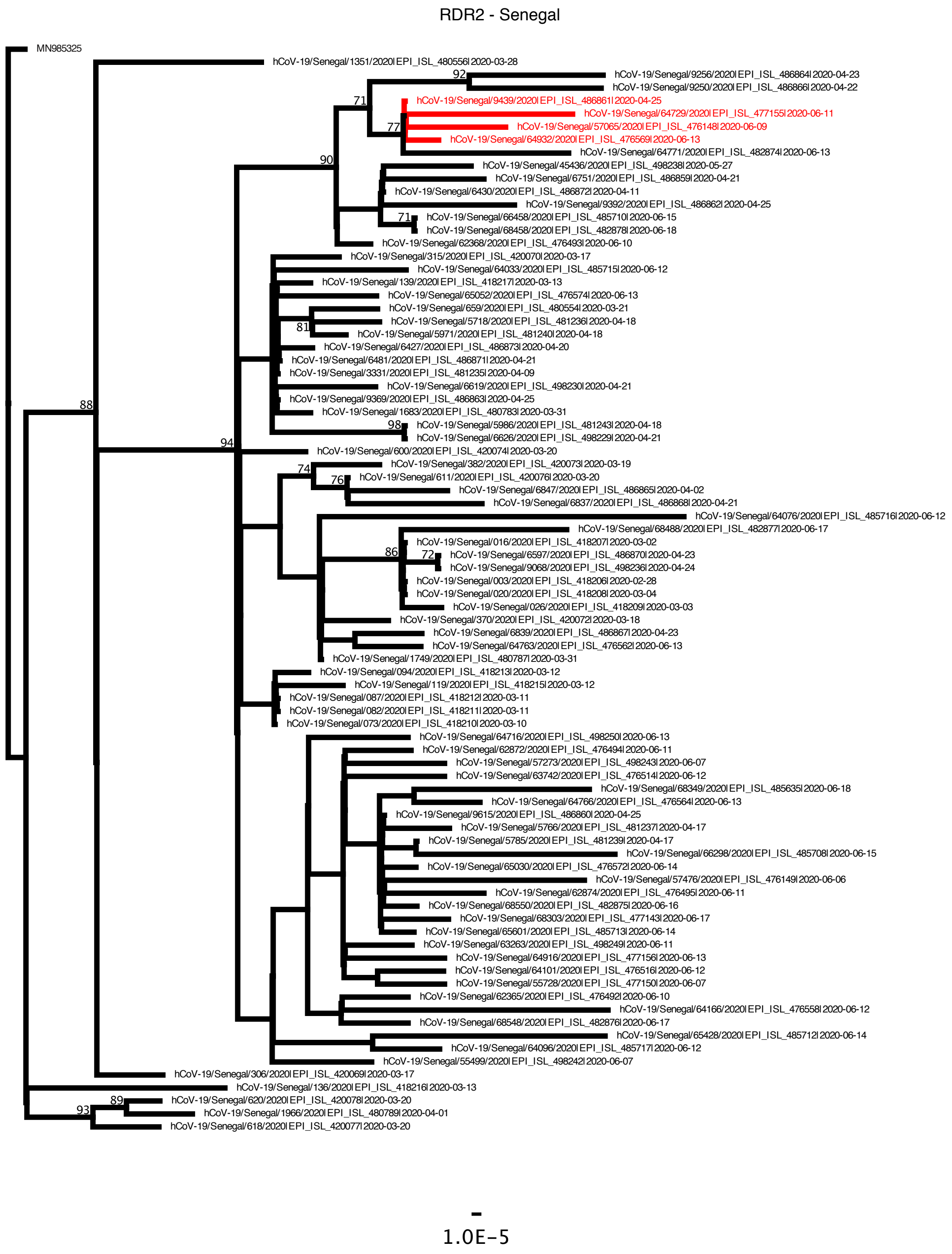


**Fig. S3**

Continued.

**E**

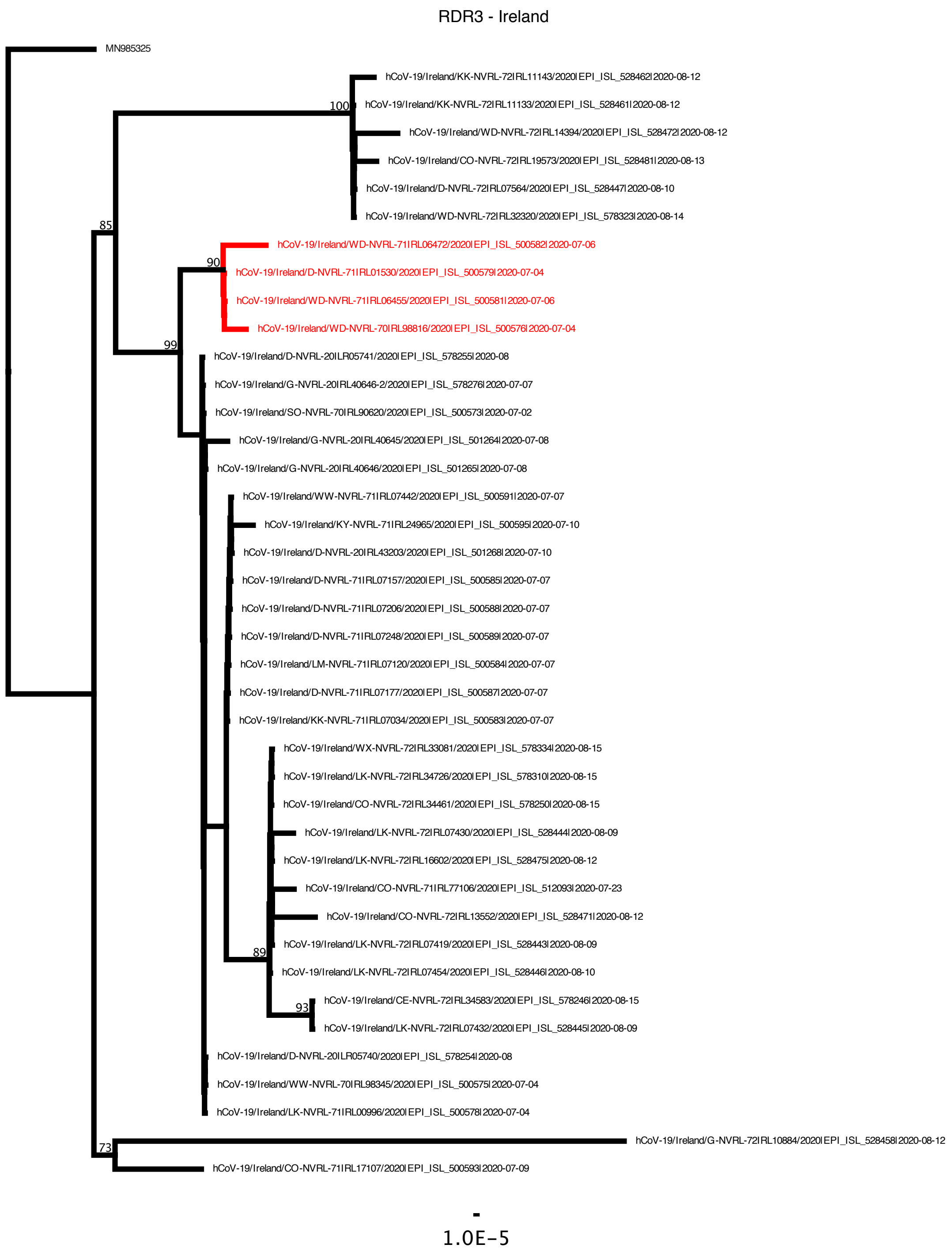


**Fig. S3**

Continued.

**F**

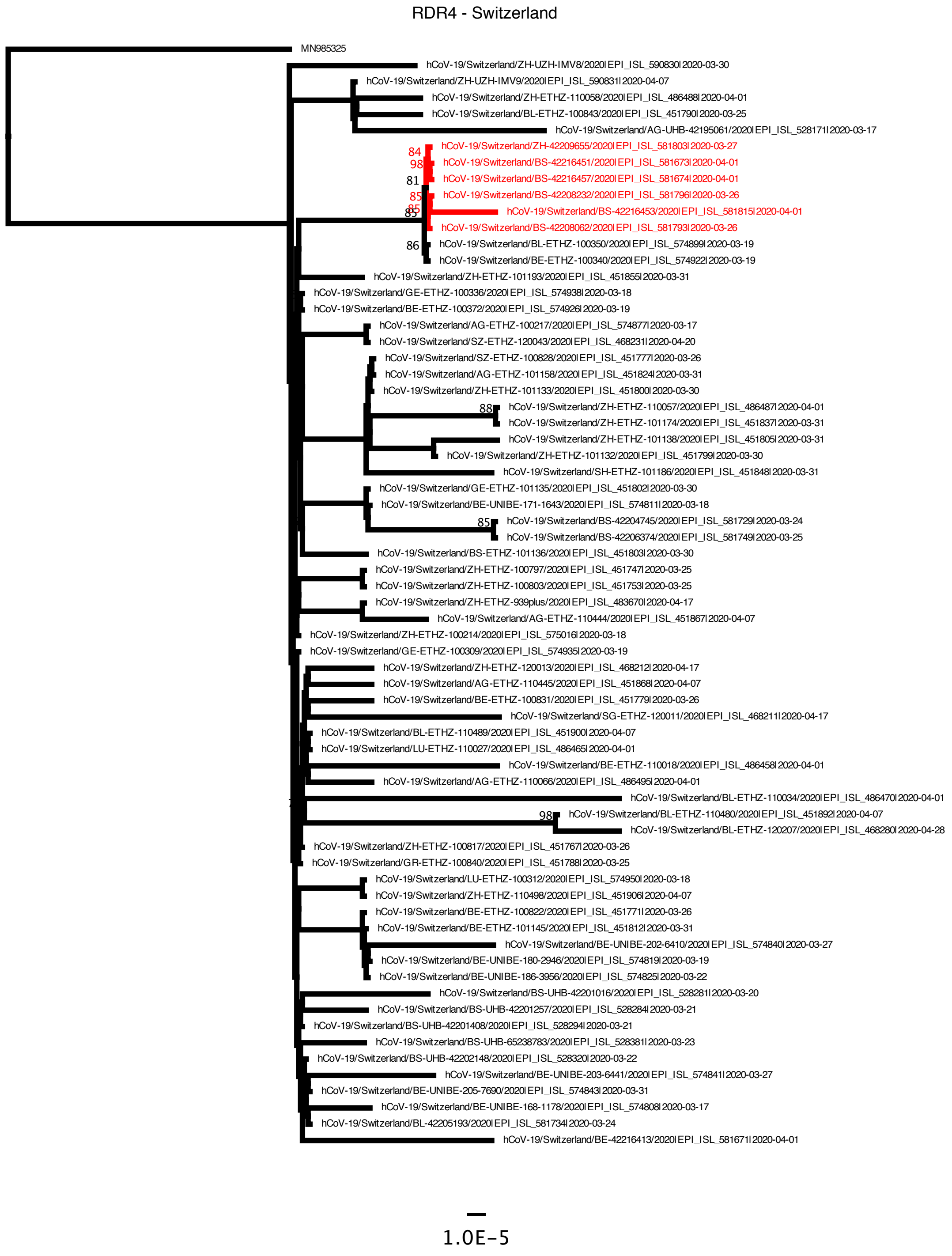


**Fig. S3**

Continued.

**Table S1: Neutralization of PLTI1 and Munich: P3 viruses using human convalescent sera**

| human convalescent serum donor | virus | serum dilution | | | | | |
|---|---|---|---|---|---|---|---|
| | | 1:25 | 1:50 | 1:100 | 1:150 | 1:200 | 1:250 |
| 1 | PLTI1 | + | + | + | - | - | - |
| | Munich: P3 | + | + | + | - | - | - |
| 2 | PLTI1 | + | + | + | - | - | - |
| | Munich: P3 | + | + | + | - | - | - |
| 3 | PLTI1 | + | + | - | - | - | - |
| | Munich: P3 | + | + | - | - | - | - |
| 4 | PLTI1 | + | + | + | - | - | - |
| | Munich: P3 | + | + | + | - | - | - |
| | | serum dilution | | | | | |
| | | 1:100 | 1:150 | 1:200 | 1:250 | 1:300 | 1:350 |
| 5 | PLTI1 | + | - | - | - | - | - |
| | Munich: P3 | + | + | + | - | - | - |
| 6 | PLTI1 | + | + | + | + | - | - |
| | Munich: P3 | + | + | + | + | - | - |
| | | serum dilution | | | | | |
| | | 1:200 | 1:300 | 1:400 | 1:500 | 1:600 | 1:700 |
| 7 | PLTI1 | + | + | + | + | - | - |
| | Munich: P3 | + | + | + | + | + | - |
| 8 | PLTI1 | + | + | + | + | + | - |
| | Munich: P3 | + | + | + | + | + | - |

+ Neutralization (by PRNT 80)
- No Neutralization (by PRNT 80)

**Table S1.**

**Additional Data: Table S2 (separate file)**
GISAID Acknowledgement Tables
